# Inverse-scattering of absorptive samples via beam propagation

**DOI:** 10.64898/2026.04.21.719764

**Authors:** Peter Wagenaar, Jeongsoo Kim, Mary E. Swartz, Johann K. Eberhart, Shwetadwip Chowdhury

## Abstract

Inverse-scattering methods enable label-free, quantitative visualization of a sample’s three-dimensional (3D) refractive index (RI), providing intrinsic and volumetric morphological contrast without exogenous labels. This is achieved by developing computational frameworks that reconstruct the sample’s 3D RI from a series of scattering measurements acquired under different data-capture conditions. Recent advances have demonstrated successful 3D RI reconstructions in multiple-scattering samples using angle-varying illuminations; however, these studies have primarily focused on non-absorptive samples. Here, we extend the multi-slice beam propagation (MSBP) inverse-scattering framework to reconstruct *complex-valued* RI, encompassing *both* the sample’s conventional RI (real part) and absorptivity (imaginary part). We show that reconstructing complex-valued RI makes the inverse problem ill-posed under angle-varying illumination alone, and that incorporating measurement diversity from *both* angle-varying illumination *and* sample defocus is necessary to ensure stable and accurate convergence. Experimental demonstrations were conducted on 1) dyed microsphere samples to characterize accuracy of reconstructed RI and absorptivity; and 2) diverse absorptive scattering samples to demonstrate biological utility. These results represent an important step for label-free volumetric imaging in biological tissue, which typically exhibits *both* scattering *and* absorption.

## 1. Introduction

Optical imaging is fundamentally constrained by scattering, which can quickly degrade imaging quality in biological tissue starting at depths of just a few tens of microns. Inverse-scattering methods seek to address this challenge by computationally undoing scattering effects to reconstruct a sample’s volumetric refractive index (RI), which provides label-free quantitative morphological contrast. Traditional methods for 3D RI reconstruction have relied on *weak-scattering* approximations, such as the first Born [1] or Rytov [2], to linearize the relationship between the sample’s 3D RI and its scattering measurements [3], [4], [5]. This approach allows efficient and analytical reconstruction of the sample’s RI and has enabled a wide range of applications, including studies of cell morphology [6], [7], [8], developmental processes [9], pathology [10], [11], [12], [13], etc.

However, because such approaches rely on the weak-scattering assumption, they are generally limited to imaging *thin* biological samples with thicknesses of only a few tens of micrometers. Complex biological samples (e.g., multicellular complexes, such as organoids [14], tissue explants[15], embryos [16], etc) are often thicker and exhibit *multiple-scattering*. 3D RI reconstruction in such samples requires computational reconstruction frameworks that move beyond weak-scattering approximations, and thus often employ nonlinear, iterative strategies. Recent examples include beam propagation-based methods [17], [18] modified-Born series [19], [20], transmission matrix approaches [21], [22], physics-informed neural networks [23], [24], and Lippmann-Schwinger-based formulations [25], [26]. Using such frameworks, impressive imaging results have been obtained in various classes of multicellular scattering samples, such as organoids [14], [27] and small organisms (e.g., *C. elegans*, tardigrades, mouse and zebrafish embryos) [28], [29], [30].

Notably, all the inverse-scattering demonstrations described above neglected absorptive effects, as they focused on samples that were scattering but largely non-absorptive. This leaves an important application gap, since biological tissues are generally *both* multiple-scattering *and* absorptive. Moreover, absorption often carries critical information about tissue composition, function, and physiological state that cannot be obtained from morphology alone. For example, absorbers endogenous to biological tissue, such as hemoglobin, melanin, lipids, and metabolic chromophores (e.g., cytochromes and flavins) provide insights into oxygenation, metabolism, and vascular density, which are not easily captured by RI alone [31], [32], [33]. Other important, more application-specific examples include retinal tissues, which contain photoreceptors with well-characterized wavelength-dependent absorption. Obtaining absorption information from these tissues can provide insight into the visual transduction processes [34], [35]. Plant tissues also contain strongly absorptive components, such as chlorophyll, carotenoid, and anthocyanin pigments [36], [37]. Absorption-based information of such tissues can be used to estimate biochemical traits (e.g., nitrogen content, specific leaf area, water status) [38], [39], [40], [41] and even physiological status (e.g., nutrient deficiency, senescence, etc) [41], [42], [43]. Thus, reconstructing 3D absorption in multiple-scattering samples would have broad and impactful applications across the sciences.

Importantly, though absorption and RI are often treated as separate properties, they are fundamentally linked. Specifically, absorptivity is determined by the imaginary component of complex-valued RI. *Thus, inverse-scattering methods that reconstruct complex-valued RI should, in principle, inherently also recover the sample’s 3D absorptivity*. To the best of our knowledge, this has not yet been demonstrated in practice in multiple-scattering samples.

*In this work, we demonstrate for the first time an inverse-scattering methodology that reconstructs 3D <u>complex-valued</u> RI in thick multiple-scattering tissues*. To accomplish this, we build upon our previous work with multi-slice beam propagation (MSBP), which employed a nonlinear physics-based model to reconstruct *real-valued* RI from scattering measurements captured under various illumination angles [18], [28]. In this work, we show that reconstructing complex-valued RI makes the inverse problem ill-posed with only angled-illumination measurements. To address this challenge, we introduced optical refocusing as an additional source of measurement diversity. We validate that MSBP achieves correct convergence with this approach, by reconstructing 3D absorptivity and RI in various calibrated, dyed microspheres. We also conduct demonstrations in various multiple-scattering bio-samples containing endogenous absorbers. *To the best of our knowledge, this work presents the first joint reconstructions of 3D RI and absorptivity in samples that exhibit <u>both</u> multiple-scattering and absorption*. This represents an important step toward label-free volumetric imaging of scattering and absorptive biological tissues.

## 2. Computational framework

### 2.1 Multi-slice beam propagation for complex-valued RI reconstruction

We begin by introducing our inverse-scattering framework to reconstruct 3D absorptivity (jointly alongside conventional 3D RI). As described above, this is achieved by extending the formulation for conventional MSBP towards reconstructing *complex-valued* RI.

Conventional MSBP decomposes a 3D scattering sample into a sequence of infinitesimally thin layers, each represented by a 2D map of *real-valued* RI [1]. The transmittance function corresponding to each layer thus acts to *phase-only* modulate the optical field incident to that layer. The sample’s overall scattering characteristics are approximated by accumulating the diffraction effects that occur as light propagates layer-by-layer through the sample.

Our extension to this model keeps a similar intuition, with the key difference that the RI is now allowed to be complex-valued. The real component corresponds to the conventional RI term while the imaginary term encodes absorptivity. As a result, the transmittance function of each layer can be decomposed into a phase-only modulation term, as before, and an additional real-valued exponential-decay term, which accounts for absorption. Thus, as light propagates layer-by-layer through the sample’s complex-valued RI, it undergoes both phase-modulation and absorption (see **Figure 1**). We describe this mathematically.

**Figure 1.**
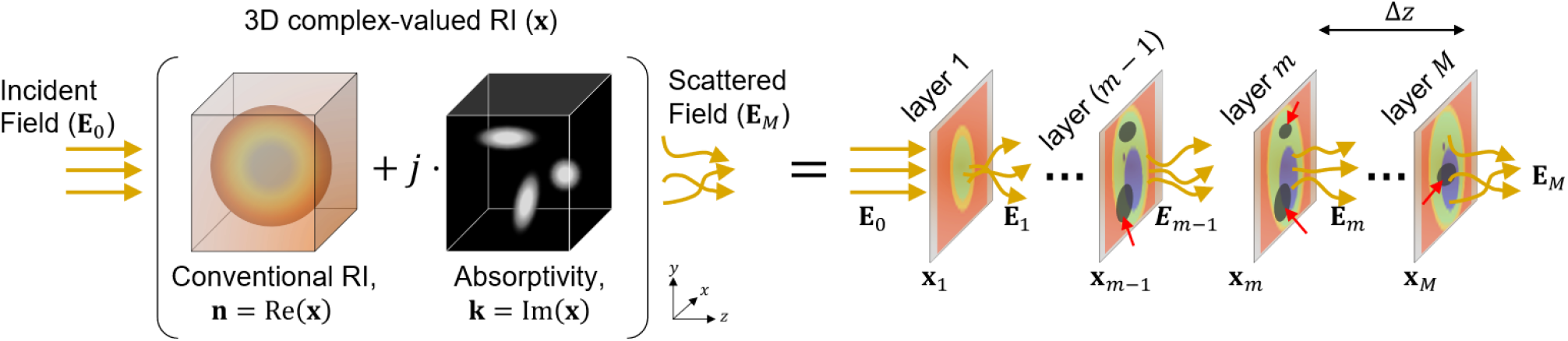
Illustration of MSBP forward model for absorptive scattering samples. Samples are represented as 3D complex-valued RI volumes, comprised of terms for conventional RI and absorptivity. In the layer-by-layer depiction, color represents conventional RI, while dark regions (marked by red arrows) highlight areas of absorption.

Consider a 3D scattering sample decomposed into *M* layers separated by a distance Δ*z*. In MSBP, light propagation through the sample is modeled as an alternating sequence of free-space propagation and layer-wise modulation steps. We mathematically describe this process below with vector/matrix variables, keeping similar notation as that introduced by Kamilov *et al*. [17]:

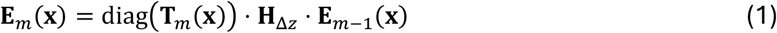

Here, **x** = **n** + *j***K** is a vector representing the sample’s 3D complex-valued RI. **n** and **K** are the real and imaginary components of **x**, representing (conventional) RI and absorptivity, respectively. **E**_*m−1*_ represents the field exiting the (*m* − 1)-th layer. We matrix-multiply this field with matrix operator **H**_Δ*z*_, which represents optically propagating **E**_*m−1*_ a distance Δ*z* to the *m*-th layer. It is then spatially modulated by the *m*-th layer’s transmittance function, 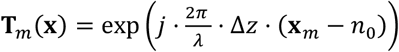, where *n*_0_ denotes the homogenous, eal-valued RI of the surrounding, non-absorptive media. **x**_*m*_ = **n**_*m*_ + *j***K**_*m*_ represents the sample’s complex-valued RI restricted to the *m*-th layer (and zero elsewhere). Decomposing the full 3D sample into its constituent layers is equivalent to the summation **x** = **x**_*1*_ + **x**_*2*_ + ⋯ **x**_*M*_.

Modulation of the field propagated to the *m*-th layer (i.e., **H**_Δ*z*_ ⋅ **E**_*m−1*_) by that layer’s transmittance function corresponds to a spatially point-wise multiplication. This is represented in Eq. (1) as a matrix-multiplication by the diagonal matrix diag(**T**_*m*_). The resulting field, denoted **E**_*m*_, represents the optical field exiting the *m*-th sample layer. Starting from the illumination field incident to the sample, **E**_0_, Eq. (1) repeated layer-by-layer, yielding the final field exiting the sample, **E**_*M*_.

Notably, by decomposing **x**_*m*_ into its real and imaginary components, **T**_*m*_ can be re-expressed to explicitly capture its phase-modulation and absorptive effects:

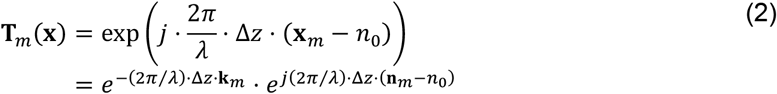

From the expanded form of Eq. (2), we see that **T**_*m*_ comprises a purely real-valued exponential term accounting for absorption (first term) and the conventional phase-only modulation term (second term). Thus, Eq. (2) describes how light is both phase-modulated *and* absorbed at each sample layer as it propagates through the volume. Notably, in the case where there is no absorption (i.e., **K = 0**), Eqs. (1) and (2) naturally simplify to the conventional MSBP model [17], [18].

In practice, measurements are acquired with the camera focused at a plane within the 3D sample volume. Thus, the MSBP *forward model* *G* describing the field at the measurement plane can be written as:

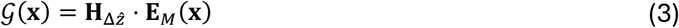

Recall that **E**_*M*_ denotes the field exiting the final (i.e., *M*-th) sample layer. Matrix multiplication by 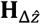 corresponds to back-propagating this field by a distance 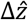 to the in-focus imaging plane within the sample. Notably, the optical propagation operator **H** in Eqs. (1) and (3) was implemented in practice using the angular spectrum method in the spatial-frequency domain, while also simultaneously applying spatial-frequency filtering to enforce a diffraction-limited pupil function [44].

### 3.2 Inverse problem

Thus, for a given volumetric RI distribution **x**, the forward model *G*(**x**) in Eq. (3) predicts the scattered field at the measurement plane. In practice, our goal is to recover **x** from the measurements which constitutes the *inverse-problem*. Because RI is a three-dimensional quantity while each measurement is inherently two-dimensional, this inverse problem is fundamentally ill-posed. Robust reconstruction therefore requires acquiring a sequence of measurements with sufficient diversity to adequately constrain the inverse problem.

To accomplish this, we acquired multiple scattering measurements of the sample being illuminated with various angled plane-waves as well as being defocused to multiple positions. Consider the sample illuminated with the *ℓ*-th plane-wave and defocused to the *p*-th position. In this case, we can generalize the MSBP forward model defined in Eq. (3) as *G*^(*ℓ,p*)^(**x**), where *ℓ* and *p* index the illumination angle and sample-defocus distance, respectively. Notably, the *p*-th sample-defocus distance can be naturally incorporated by generalizing 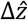 in Eq. (3) as 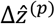.

The inverse-scattering framework for reconstructing the sample’s complex-valued RI follows naturally. We formulate this problem as a least-squares optimization that searches for the 3D RI distribution that minimizes the discrepancy across all measurements between the measured scattering *amplitudes* (i.e., square-root of raw intensity) and the *amplitude* predictions made by the MSBP forward model in Eq. (3):

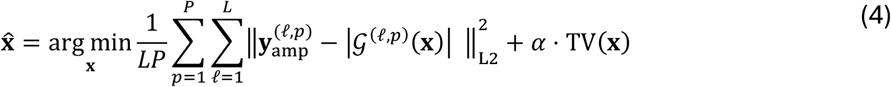

Here, 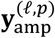 denotes the measured scattering *amplitude* corresponding to the *ℓ*-th plane-wave illumination and the *p*-th sample-defocus position. *L* and *P* denote the total number of illumination angles and sample-defocus positions, respectively. ‖ … ‖_*L2*_ denotes the L2-norm operator. The term *TV(…)* denotes the 3D total-variation regularizer, which stabilizes the iterative optimization process from noise and poor conditioning, while also mitigating missing-cone artifacts [45]. *α* sets the strength of regularization. In practice, TV regularization is implemented via proximal operator; details are provided in **Supplementary Note S1**.

The inverse problem formulated in Eq. (4) is structurally similar to those presented in prior works [17], [18], [30]. A key distinction, however, lies in its implementation. In particular, solving Eq. (4) requires computing the gradient of the least-squares objective, which in previous formulations was derived under the assumption that the sample RI **x** was purely real-valued. In this work, **x** is explicitly complex-valued, and the least-squares term in Eq. (4) involves field amplitude that contains explicit dependence on the complex conjugate of **x**. As a result, Eq. (4) defines a complex-valued, non-holomorphic optimization problem. Accordingly, we re-derived the gradient of the least-squares term using Wirtinger calculus; full derivations are provided in **Supplementary Note S1**.

As we show below, our empirical results demonstrate that reconstructing complex-valued RI from angle-varying illumination alone leads to an ill-posed inverse-problem, producing RI reconstructions with pronounced artifacts. For this reason, we increased measurement diversity by combining angular illumination with sample defocus. We observed that this strategy provided additional constraints that enabled accurate reconstruction of the sample’s complex-valued 3D RI.

## 3. Experimental Methods

### 3.1 Optical design

We show in **Figure 2** the schematic of our optical system, implemented as an off-axis interferometric setup with angle-scanning illumination. Coherent illumination was provided by three laser diodes operating at blue, green and red wavelengths (Thorlabs, CPS450, *λ*=450nm; Thorlabs, CPS532, *λ*=532 nm; Thorlabs, CPS635, *λ*=635 nm). The diodes were used independently, with each source fiber-coupled into the system one at a time. A pair of lenses then expanded and collimated the output beam to create a plane wave with a gaussian intensity profile.

**Figure 2.**
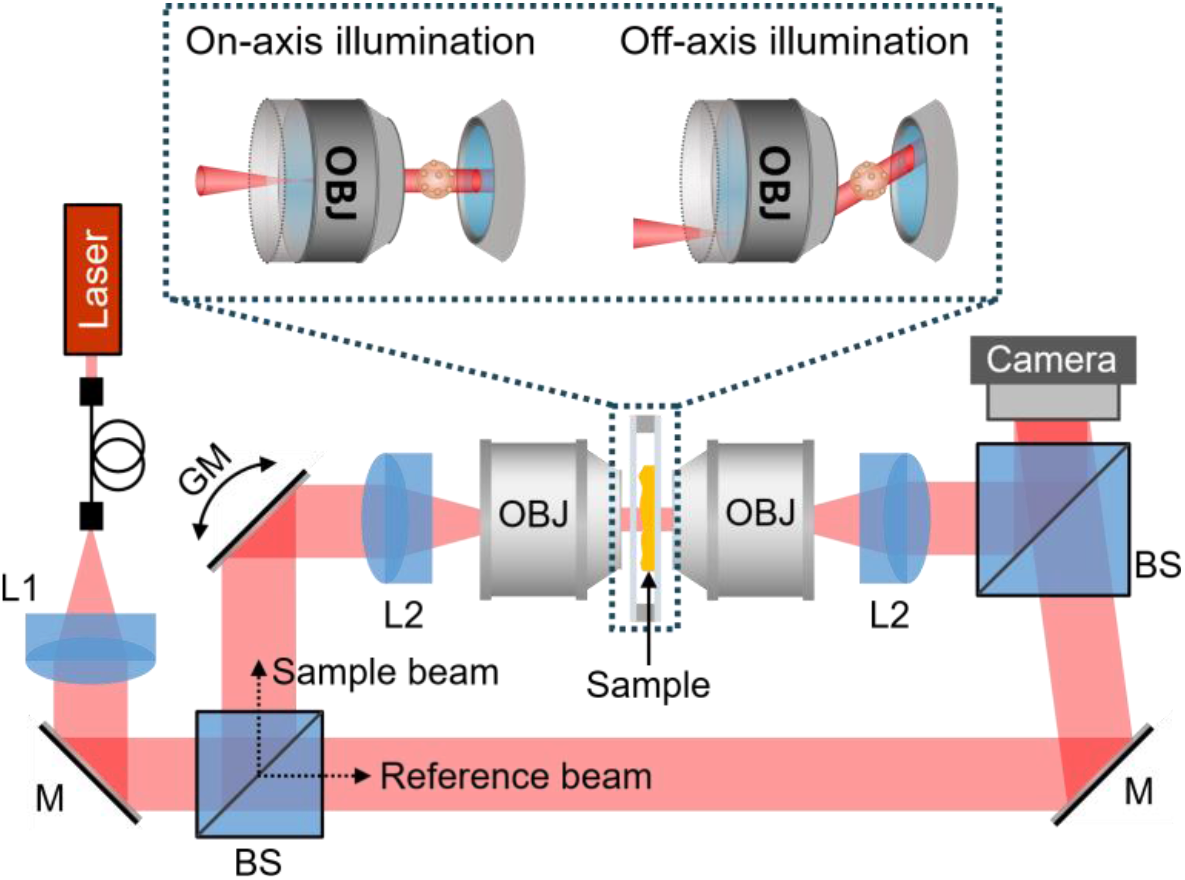
Optical schematic of angle-scanning, off-axis interferometric imaging system. For illustration simplicity, only one laser source is shown. First beam splitter (BS) splits light into sample and reference beams. Sample beam is directed to a galvo mirror (GM), which steers the illumination beam onto the sample, through 4f system L2 and OBJ. Scattered light from the sample is imaged through an identical 4f system onto the camera. Reference beam is directed to the camera at an off-axis angle from the sample beam.

A 50/50 beam splitter (BS) (Thorlabs, CCM1-BS013) split the incoming light into sample and reference beams that propagates through two separate arms of the system. In the sample arm, light traveled to a 2D scanning galvanometer (GM) (Thorlabs, GVS212), which steered the beam to illuminate the sample across a range of incident angles. The GM and camera (ToupTek, IUA20000KMA, Pixel Pitch = 2.4 um) were in conjugate imaging planes. From the GM, the beam was illuminated into the sample volume by a 4f system consisting of a tube lens (Thorlabs, ACT508-200-A-ML, f=200mm) and an objective lens (OBJ). The resulting scattered light from the sample was then imaged with another objective and tube lens to the camera. For characterization experiments with microspheres, we used identical high-resolution objective lenses for both illumination and imaging (Nikon 60x, 1.42 NA oil) to maximize resolution. For imaging thicker biological samples, we used objectives with longer working distance (illumination: Olympus, 20x, 1.0 NA water; imaging: Nikon, 16X, 0.8 NA water).

The reference beam was directed by a mirror (M) onto the camera at an off-axis angle relative to the sample beam, creating interference fringes at the camera’s imaging plane. From this interferogram, standard principles from off-axis holography were used to measure the complex-valued field at the camera plane [46]. The position and tilt of the reference beam were aligned so that the reference and sample beams spatially overlap, while in Fourier space the central DC component was sufficiently separated from the +/-first order components, which contained the relevant complex-valued information of the sample field.

### 3.2 Data acquisition

Interferometric measurements were acquired as the sample was sequentially illuminated with multiple angled plane waves. Illumination angles were arranged along a spiral trajectory spanning the full extent of the imaging objective lens’s pupil area. Acquisition parameters are described for each acquired dataset in **Supplementary Note S4**.

From each interferometric measurement, complex-valued field (amplitude and phase) was recovered by computationally isolating the first order diffraction term of the interferogram in the spatial-frequency domain [46]. This process was repeated after translating the sample out of the field of view in order to collect background field measurements. The sample fields were then normalized with the corresponding background fields via complex-valued division. Finally, the amplitude-only “measurements” used as *Y* in Eq. (4) above were synthetically generated by simply taking the magnitude of the normalized field measurements. Notably, the illumination angles for each measurement were computationally recovered with high precision directly from the non-normalized background field measurements by locating the DC peak in the spatial-frequency domain.

Lateral and axial resolution in angle-scanning RI-tomography are given by *d*_*xy*_ = *λ*/2NA and 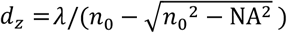, respectively, where we recall that *n*_0_ denotes the RI of the surrounding medium [47]. For the microsphere characterization experiments, a 1.42 NA oil-immersion (*n*_0_ = 1.515) objective was used, yielding lateral resolutions of *d*_*xy*_ = 160 nm, 190 nm, and 220 nm, and axial resolutions of *d*_*z*_= 450 nm, 530 nm, and 640 nm, for wavelengths *λ* = 450 nm, 532 nm, and 635 nm, respectively. For the biological imaging experiments, a 0.8 NA water-dipping objective (*n*_0_ = 1.33) was used at a single wavelength of *λ* = 635 nm, giving a lateral resolution of *d*_*xy*_ = 400 nm and an axial resolution of *d*_*z*_= 2.4 µm.

### 3.4 Digital defocusing to increase measurement diversity

As described earlier, solving Eq. (4) involves navigating a non-holomorphic, complex-valued solution space. This space is generally highly nonconvex, which poses significant challenges for accurate convergence. Our experimental results (discussed below) indicated that angled illuminations alone were not sufficient to achieve accurate convergence for complex-valued RI reconstruction. Thus, we increased measurement diversity by also incorporating sample defocusing.

Because we collect complex-valued *field* measurements (**Y**_*field*_), we could implement sample-defocusing *fully computationally* by numerically propagating the measured field across varying distances, i.e., 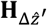, **y**_*field*_, where 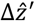 corresponded to the defocus distance. This allowed us to defocus the sample *without physical sample movement or additional acquisition time*. The amplitude component of each digitally defocused field was then used as a “measurement” **Y**_*amp*_ within the inverse problem in Eq. (4).

In summary, we captured scattering field measurements for each sample at *L* illumination angles. Each angular field measurement was then numerically propagated across *P* defocus planes, after which the amplitude component of the field was extracted. Thus, each unique combination of angle and defocus can be represented as 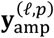 in Eq. (4), where *ℓ* and *p* index the illumination angles and defocus planes, respectively. Information of *L* and *P* for each sample, as well as specific sample-defocus positions, is provided in **Supplementary Note S4**. Importantly, 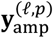 and *G*^(*ℓ,p*)^(**x**) in Eq. (4) both index the same physical sample-defocus position.

A key insight of this approach is that defocusing each angular measurement indirectly encodes sample-specific phase information despite each individual measurement term *Y* in the solver being purely a real-valued amplitude. *We intentionally omit direct phase information within the inverse solver*; we have recently shown that in multiple-scattering samples, even real-valued RI reconstructions derived directly from complex-valued fields are prone to severe artifacts [30].

## 4. Results

### 4.1 Quantitative characterization

To validate the quantitative imaging capabilities of our system, we first reconstruct the complex-valued RI of absorptive polystyrene microspheres (PolySciences, PolyBead Polystyrene 10um diameter Microspheres; Blue: 18138, Yellow: 18337, Black: 24294, and Violet: 18139) immersed in nonabsorptive, index-matching oil (Cargille Series A 1.572), and compare them with the microspheres’ “ground-truth” RI values. Importantly, while the RI of *undyed* polystyrene is well characterized across a broad wavelength range, these reference curves assume a purely real-valued RI and do not account for absorption, which is known to modify the real component of RI [48].

Instead, to obtain ground-truth complex-valued RI values for the absorptive microspheres, we analyzed two-dimensional phase and amplitude maps extracted from field measurements acquired at normal incidence of the illumination beam. Because a monolayer of microspheres well index-matched to the surrounding medium is weakly scattering, the measured 2D amplitude and phase maps corresponded to projections of the imaginary (absorptivity) and real (conventional RI) components of the microsphere’s complex-valued RI, respectively. Since the microspheres were homogeneous, say with a constant complex-valued RI of *n + jk*, we can write an expression for the measured complex-valued sample field:

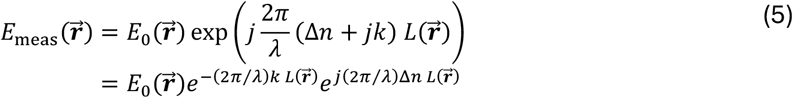

Here, *E*_*meas*_ and *E*_0_ denote the measured and incident complex-valued fields, respectively, expressed as functions of the two-dimensional *vector* of spatial coordinate 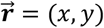. Here, *x* and *y* are *scalar* variables representing the two orthogonal cartesian directions. The quantity Δ*n* = *n* − *n*_0_ represents the RI contrast between the microsphere’s real-valued RI component (*n*) and that of the surrounding medium (*n*_0_). The term 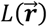 denotes the physical path length through the sample at position 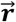. For a microsphere, 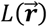 takes the form of a half-ellipse, with an extent and center peak both equal to the microsphere’s physical diameter.

To isolate the microsphere-specific contribution to the field, we normalized the measured field by the incident field, *E*_*sample*_ = *E*_*meas*_/*E*_0_. From this normalized field, the imaginary (i.e., absorptivity, *k*) and real (i.e., conventional RI, *n*) components of the microsphere’s complex-valued RI could be directly inferred from the amplitude (i.e., |*E*_*sample*_|) and phase (i.e., ∠ *E*_*sample*_), respectively:

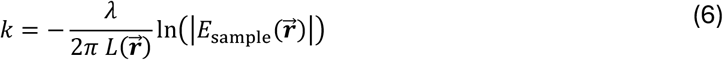

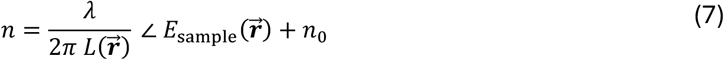

Using these relationships, we computed “ground-truth” values for *n* and *k* that provided the best fit to the measured amplitude and phase distributions across the centers of the microspheres. **Figure 3** below shows a representative fit to the measured phase and amplitude profiles of a blue-dyed microsphere with diameter 8.54 μm, recorded using red illumination (*λ* = 635 nm). We repeat this process for all colored microspheres, across blue, green, and red illuminations. Ground-truth values for absorptivity (i.e., *k*) and RI-contrast (i.e., Δ*n*) are provided in **Table S3.1** in **Supplementary Note S3**. We report Δ*n = n − n*_0_ rather than *n* because both *n* and *n*_0_ depend on wavelength, and reporting Δ*n* provides a more concise representation in the table.

**Figure 3.**
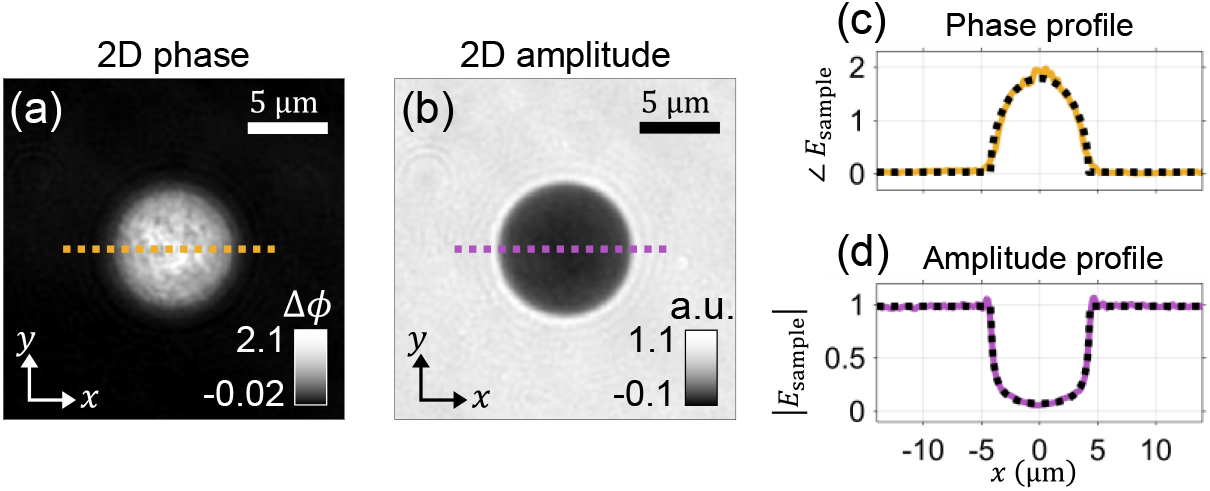
Ground-truth RI (*n*) and absorptivity (*k*) values of the calibration microspheres are obtained by fitting model profiles to those extracted from 2D **(a)** phase and **(b)** amplitude maps measured under orthogonal illumination. Line profiles taken along the dashed yellow and purple paths in (a) and (b) are shown as colored curves in **(c)** and **(d)**, respectively. The black dashed curves show the fitted phase and amplitude profiles corresponding to the inferred ground-truth values of *n* and *k*.

Next, we reconstructed the 3D RI of the dyed microspheres by solving the inverse problem in Eq. (4). **Figure 4** shows the reconstructed 3D RI-contrast for the same blue microsphere as before using: (1) the traditional MSBP formulation, which assumes a real-valued RI and uses the corresponding gradient; and (2) our extended MSBP framework for complex-valued RI reconstruction, which employs the complex-valued gradient derived in **Supplementary Note S1**.

**Figure 4.**
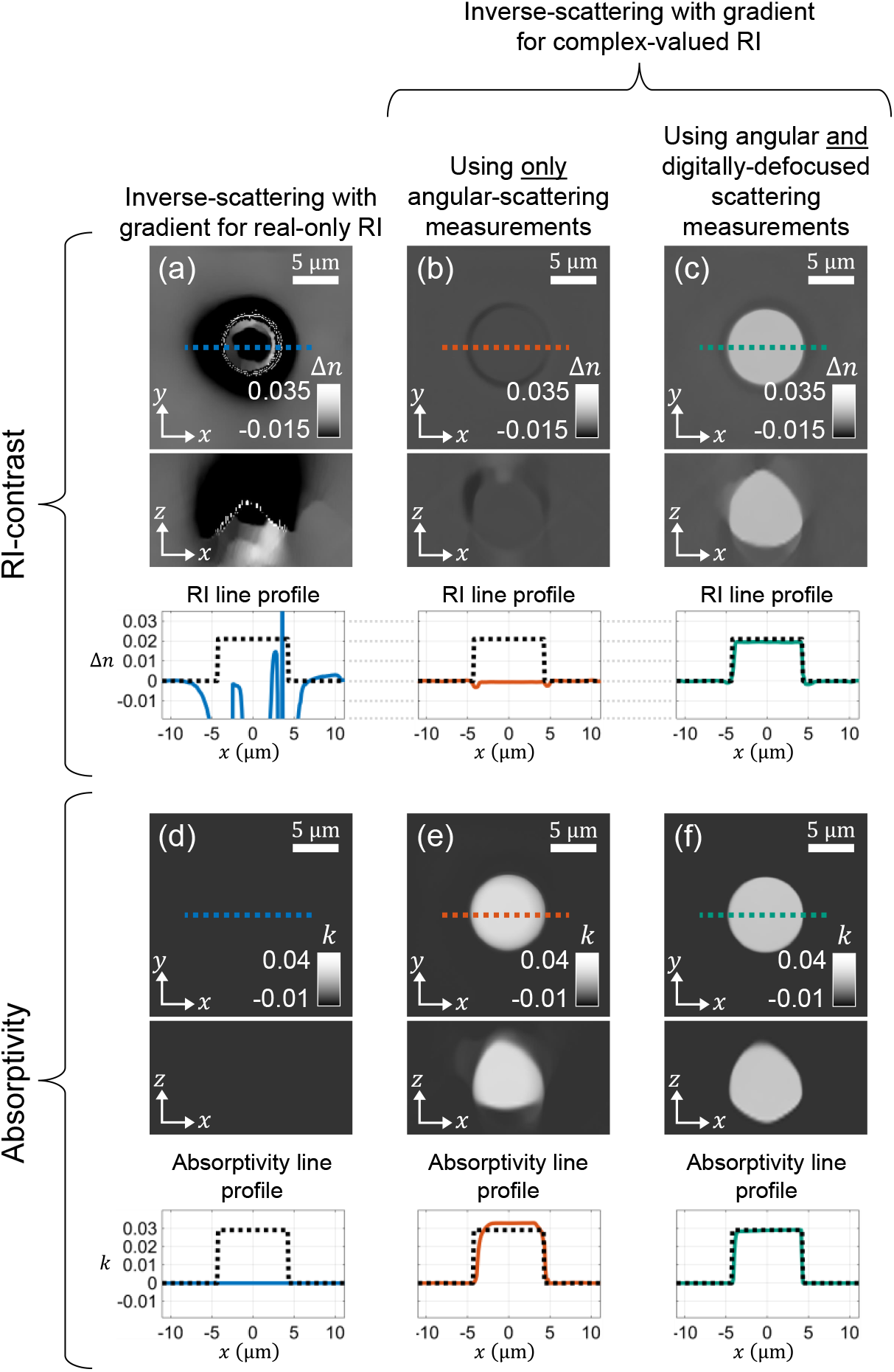
Comparison of reconstructed 3D complex-valued RI (i.e., conventional RI and absorptivity) of an absorptive (blue-dyed) microsphere. Reconstructions were obtained via MSBP-based inverse-scattering with a **(a)** gradient formulated for real-only RI and using data from angular scattering measurements, as well as a gradient formulated for complex-valued RI and using **(b)** only angular scattering measurements and **(c)** both angular *and* defocused scattering measurements. Line profiles extracted along the dashed lines in (a – f) are shown to assess quantitative accuracy across all reconstruction scenarios. Black dashed curves in the plots denote the “ground-truth” RI and absorptivity values computed from measured 2D fields, as described previously in Fig. 2.

When the inverse problem was solved using the gradient derived for real-valued RI reconstruction, the resulting RI distribution was severely inaccurate and exhibited sharp discontinuous spikes of RI values surrounded by regions of extremely low RI. **Figure 4(a)** demonstrates this failed reconstruction specifically in the case when only angular scattering measurements were used – however, similarly failed results were obtained even in the case when digitally defocused measurements were included (see **Supplementary Note S2**). This outcome was expected, as the assumption of a purely real RI was fundamentally inconsistent with absorptive microspheres.

In contrast, using the gradient derived for complex-valued RI reconstruction led to immediate and substantial improvements. As shown in **Figure 4(b,e)**, reconstructions using only angular scattering measurements revealed a clear silhouette of the microsphere in both RI and absorptivity, and were qualitatively more representative of the ground-truth than the reconstruction in **Figure 4(a)**. However, line profiles through the center of the volume indicated that quantitative accuracy remained poor for the reconstructed RI. In contrast, when the reconstruction was repeated using *both* angular and digitally defocused measurements (**Figure 4(c,f)**), both the 3D reconstructed RI and absorptivity were of high quality and high quantitative accuracy.

Results shown in **Figure 4** correspond to a blue-dyed microsphere illuminated with red light (*λ*=635 nm), for which the sample exhibited strongest absorption. Line-profile analysis yielded RI and absorptivity errors of *n*_*ϵ*_ < 9.3 × 10^−4^ and *k*_*ϵ*_ < 1.9 × 10^−3^, respectively, competitive with accuracies reported by other RI imaging methods [49], [50]. We repeated this analysis across all dyed microspheres and illumination wavelengths and summarized the values in **Table S3.1** in **Supplementary Note S3**. Notably, reconstruction accuracy was observed to be inversely correlated with sample absorptivity. We speculate that this was due to the reduced signal-to-noise ratio (SNR) associated with higher absorption. Even after adjusting the regularization strength *α* in Eq. (4) to compensate for SNR, reconstruction accuracy still showed modest degradation with higher absorption. For example, a yellow-dyed microsphere was substantially less absorptive under red illumination and achieved *n*_*ϵ*_ < 1 × 10^−5^ and *k*_*ϵ*_ < 1 × 10^−5^, more than an order of magnitude *lower* than the errors obtained for the blue-dyed microsphere under the same illumination conditions.

### 4.3 Biological weak-scattering sample

We next apply our technique to a moderately scattering yet strongly absorptive biological sample. Specifically, we image *Micrasterias Radiata* (MR) algae, which exhibit strong chlorophyll-related absorption in the red and blue regions of the visible spectrum. This is evident in **Figure 5(a)**, which shows a brightfield image of an alga acquired under red light, revealing extensive absorptive regions throughout the specimen. We use our MSBP inverse-scattering framework to reconstruct this sample’s 3D complex-valued RI.

**Figure 5.**
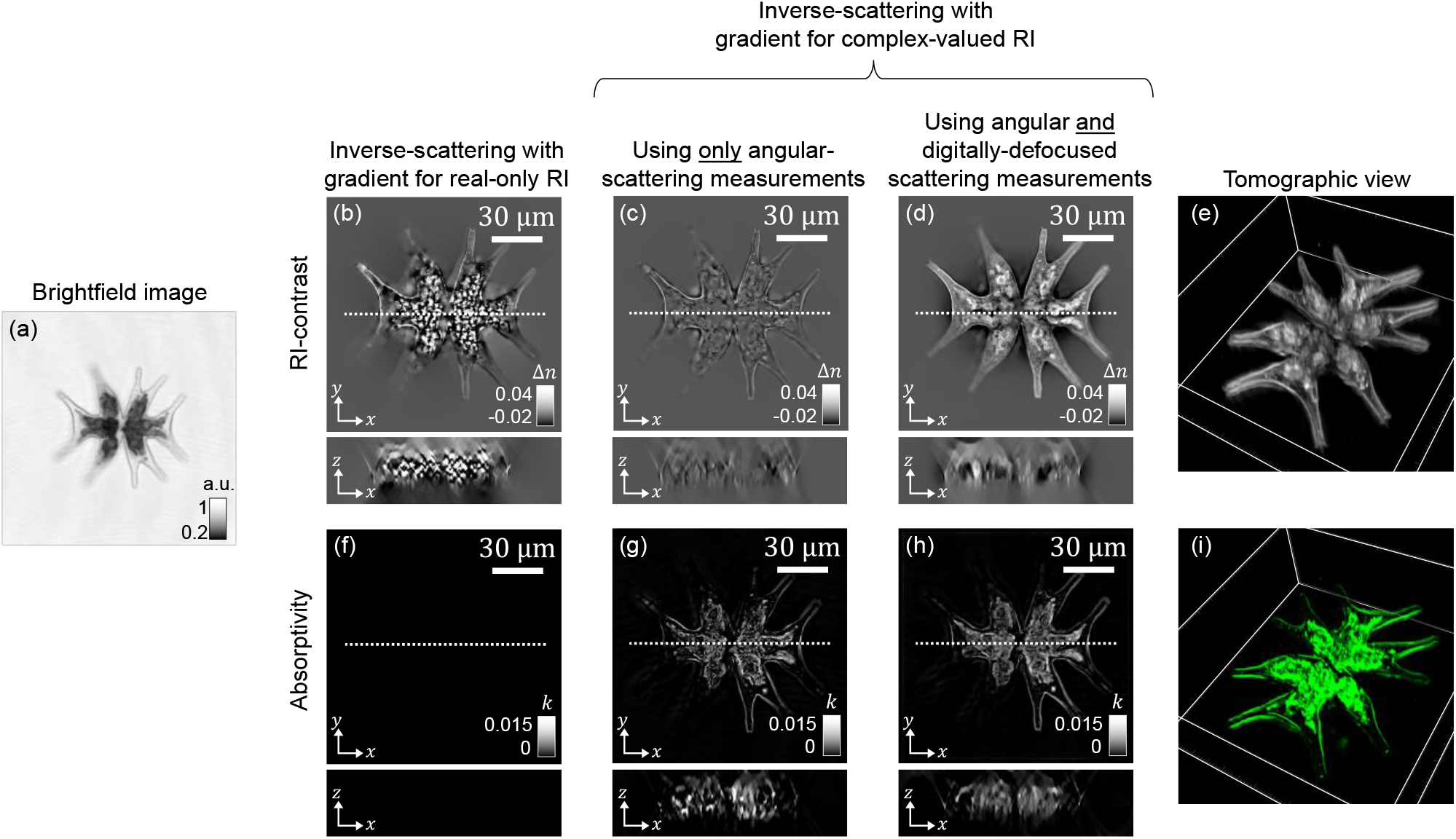
**(a)** Brightfield image of *Micrasterias radiata* under red illumination. Chlorophyll within the cell absorbs red light, producing darker regions corresponding to chlorophyll-rich structures. x-y and x-z cross-sections through the center of the reconstructed 3D RI are shown, where RI reconstructions are obtained using MSBP-based inverse scattering with **(b)** gradient for real-only RI and using angular scattering measurements, **(c)** gradient for complex-valued RI with angular scattering measurements, and **(d)** gradient for complex-valued RI with combined angular *and* defocused scattering measurements. Panels **(f**,**g**,**h)** show the corresponding cross-sections of the reconstructed 3D absorptivity maps. **(e**,**i)** Tomographic views are shown of the reconstructed RI and absorptivity maps in (d,h), respectively.

**Figure 5(b)** shows reconstruction results obtained using the traditional implementation of MSBP, where only angular scattering measurements were used in the inverse-solver and the gradient was derived under the assumption of real-valued RI. As expected, because this assumption was strongly mismatched to the absorptive nature of the sample, pronounced reconstruction artifacts were observed. In particular, the reconstructed RI exhibited severe saturation artifacts, with values spiking to non-physiological levels especially in regions known to exhibit strong absorption based on the widefield image.

We then reconstructed 3D RI using the extended formulation MSBP presented in this work, where the gradient was derived for complex-valued RI reconstruction. **Figure 5(c,g)** shows the reconstruction results when only angular-illumination measurements were used in the inverse solver. Similar to our observations with the microspheres, this approach yielded substantial improvements over real-only reconstructions – saturation artifacts were eliminated and absorptivity information became visible. However, the reconstructed RI exhibited limited contrast, with many internal features poorly resolved. This behavior mirrored the microsphere results, where the RI component was strongly underestimated when only angular measurements were used to solve the inverse problem.

**Figure 5(d,h)** shows complex-valued RI reconstruction when *both* angular and digitally defocused measurements were used in the inverse-solver. In this case, both RI and absorptivity were reconstructed with high contrast and fidelity. The characteristic morphology of *Micrasterias radiata* was clearly resolved, such as its two mirror-image semicells joined at a narrow central isthmus, as well as the subdivision of each semicell into multiple lobes arranged in a star-like pattern. Prominent chloroplasts within each semicell are clearly visible in the absorptivity maps, exhibiting a lobed architecture that follows the overall cell geometry. The strong, wavelength-dependent absorption is attributed to photosynthetic pigments (primarily chlorophylls and carotenoids [51]) concentrated within these chloroplasts. Each chloroplast contains numerous pyrenoids, recognizable by their circular starch sheaths in the RI contrast [52], which are known to be sites of carbon fixation [53]. The rigid cell wall was also resolved, revealing fine structural features such as marginal indentations and surface patterning [54], [55], [56], [57]. Together, the reconstructions in **Figure 5(b)** highlight the pronounced morphological organization and intracellular compartmentalization characteristic of this alga.

### 4.4 Biological multiple-scattering samples

#### Succulent

We subsequently applied our technique to biological specimens characterized by both scattering and absorption, beginning with a tissue section of a common succulent. **Figure 6** below shows the MSBP-based inverse-scattering reconstruction of the sample’s 3D RI and absorptivity. The inverse-problem was solved using the gradient for complex-valued RI while leveraging measurement diversity from both angular and defocused scattering measurements. Much like the previously analyzed alga, succulents exhibited chlorophyll-related absorption, as demonstrated in the red-light brightfield image in **Figure 6(a)**. As indicated by the arrows, dark regions were primarily concentrated within the micron-scale chloroplasts as well as the parenchyma cell walls surrounding the internal water-filled vacuoles.

**Figure 6.**
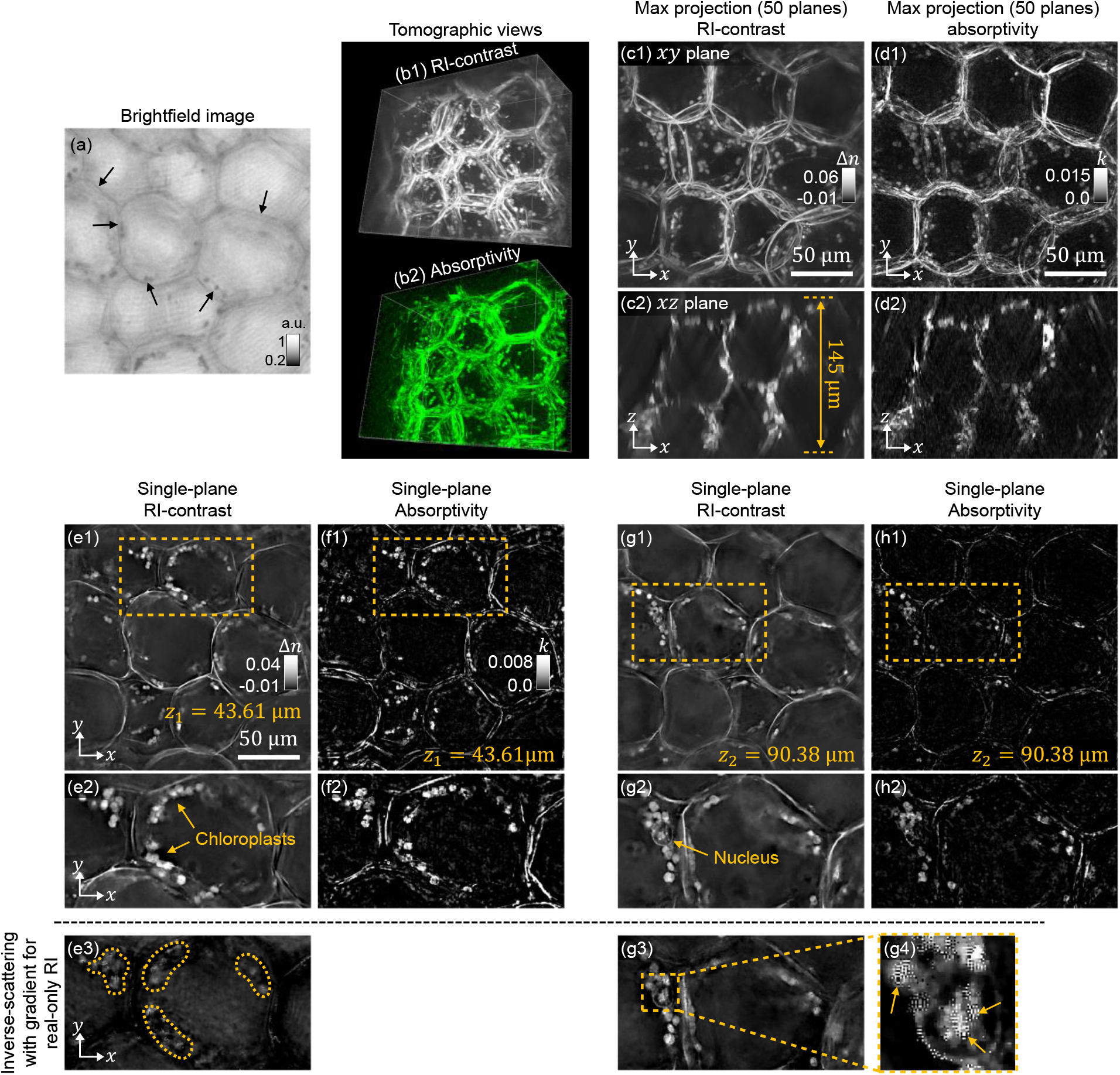
**(a)** Brightfield image of a 145-µm-thick succulent tissue section. **(b1, b2)** Tomographic renderings of the reconstructed 3D RI and absorptivity volumes obtained via MSBP-based inverse scattering using a complex-valued RI gradient and incorporating both angular and defocused scattering measurements. **(c1, c2)** Maximum x-y and x-z projections (computed over 50 planes) of the RI volume are shown, as well as **(d1, d2)** the corresponding absorptivity projections. **(e1, f1)** and **(g1, h1)** show x-y cross-sections of the RI and absorptivity volumes at *z*_*1*_ *= 43*.*61 μm* and *z*_*2*_ *= 90*.*38 μm*, respectively. **(e2, f2, g2, h2)** Magnified views are shown of the dashed yellow regions in (e1, f1, g1, h1), respectively, highlighting chloroplasts and cell boundaries with RI and absorptivity contrast. **(e3, g3)** Notably, magnified views of the same regions reconstructed using a real-only RI gradient exhibit pronounced degradation, including reduced contrast (yellow dashed regions in **(e3)**) and high-frequency saturation artifacts (indicated by yellow arrows in **(g4)**).

In **Figure 6(c1, c2)** and **Figure 6(d1, d2)**, we show x-y and x-z maximum projections (computed across 50 adjacent planes) of the reconstructed RI and absorptivity volumes, respectively. The x-z projections in **Figure 6(c2, d2)** indicate a tissue section thickness of approximately 145 µm. The absorptivity reconstruction prominently highlights the parenchyma cell walls and chloroplasts, consistent with their absorptive contrast observed in the brightfield image. Notably, these same structures also appear with high contrast in the RI maps, indicating that regions of elevated RI and absorptivity are largely spatially co-localized in this sample. **Figure 6(e1, f1)** and **Figure 6(g1, h1)** show RI and absorptivity from single x-y cross-sectional planes at axial positions *z*_1_ = 43.61 μm and *z*_*1*_ = 43.61 μm, respectively. Magnified views of the dashed yellow regions (see **Figure 6(e2, f2, g2, h2)**) reveal chloroplasts positioned along the inner periphery of the cell wall, consistent with known cellular organization [58]. **Figure 6(g2)** additionally reveals a nucleus located adjacent to the cell wall. As nuclei are predominantly phase-shifting rather than absorptive structures [55], they exhibit strong RI contrast but minimal absorptive contrast, explaining the nucleus’s clear appearance in **Figure 6(g2)** but virtually undetectable signature in **Figure 6(h2)**. Interestingly, even when using the gradient for real-only RI, the reconstructed RI appeared reasonable and partially captured some key features, such as the cell walls and the chloroplasts. This was observed likely because the succulent tissue was only weakly absorptive, as reflected by the lower contrast in its brightfield image relative to that of the strongly absorptive algal sample (see **Figure 5(a)** from before). As a result, the mismatch introduced by neglecting absorption produced less severe reconstruction artifacts than in the algal case. Nevertheless, limitations remained. For example, chloroplasts were either reconstructed with reduced contrast (**Figure 6(e3)**) or high-frequency saturation artifacts (**Figure 6(g3)**). Importantly, these artifacts were spatially correlated with regions of elevated absorptivity, mirroring the behavior observed in the algal reconstructions.

In this example, regions of elevated RI and absorptivity were largely spatially co-localized. In the next section, we reconstructed a sample in which regions of high RI and absorption are physically separate.

#### Zebrafish embryo (48 hours-post-fertilization)

Lastly, we demonstrate 3D RI reconstruction of the tail region of a 48 hours-post-fertilization (hpf) zebrafish embryo (see **Supplementary Note S5)**, a widely used multicellular system that provides insight into embryogenesis and organ formation.. However, imaging the zebrafish embryo at subcellular resolution is often challenging due to its complex architecture, which induces substantial multiple light scattering. In prior work, we applied MSBP to image a 24 hpf zebrafish embryo and successfully reconstructed rich structural detail using label-free RI contrast. At that stage, the embryo is largely non-absorptive, so assuming a real-valued RI was sufficient. In contrast, by 48 hpf, melanophore development introduces significant optical absorption. As a result, the real-only RI assumption is no longer valid, necessitating reconstruction of complex-valued RI.

**Figure 7** presents reconstruction results obtained using our proposed framework for complex-valued RI recovery using measurements combining angular illumination and sample defocus. To illustrate the global structure of the embryo we first show x-y and x-z maximum-intensity projections of both the reconstructed absorptivity and RI volumes. **Figure 7(a1, a2)** display maximum projections of absorptivity in the x-y and x-z planes. Although the embryo remained largely transparent, sparse regions of strong absorption were evident that extended across the whole embryo. The zoomed-in view in **Figure 7(a3)** reveals that these high-absorptivity regions corresponded to melanophores, identifiable by their morphology and consistency with prior reports [59], [60], [61].

**Figure 7.**
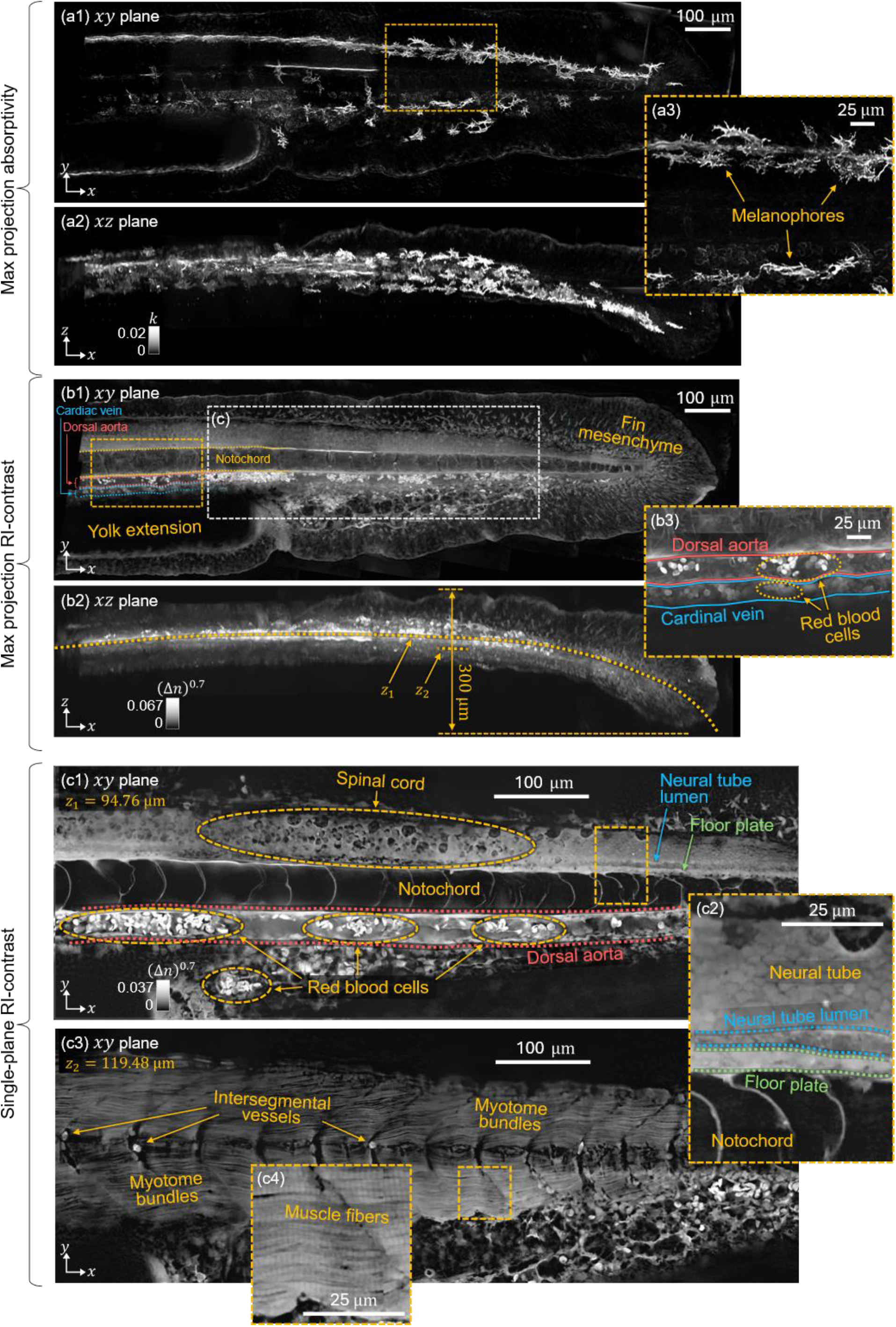
**(a1, a2)** Maximum-intensity projections of reconstructed absorptivity in the x-y and x-z planes, with the (a3) zoomed-in view identifying melanophores as the dominant absorptive feature. **(b1, b2)** Maximum-intensity projections of reconstructed RI in the x-y and x-z planes depicts the overall tail morphology, clearly visualizing the cardinal vein, dorsal aorta, and notochord. **(b3)** Zoomed-in view indicates the red blood cells visualized in the vasculature. **(c1)** Lateral cross-section at *z*_1_ = 94.76 μm resolves the notochord, spinal cord, dorsal aorta, and red blood cells, as well as the **(c2)** the neural tube, neural tube lumen, and floor plate in the zoomed-in view. **(c3)** Lateral cross-section at *z*_2_ = 119.48 μm reveals segmented myotome bundles and intersegmental vessels. **(c4)** Zoomed-in view show the individual striated muscle fibers.

**Figure 7(b1, b2)** shows maximum projections of the reconstructed RI volume. The overall morphology of a 48 hpf zebrafish tail was clearly resolved, including the cardiac vein, dorsal aorta, notochord, and yolk extension. The fin mesenchyme was also clearly visualized, within which the developing fin rays were readily distinguishable. **Figure 7(b3)** shows a zoomed-in view of the region indicated by the yellow dashed rectangle in **Figure 7(b1)**, revealing distinct red blood cells located within the dorsal aorta and cardinal vein. Interestingly, red blood cells in the dorsal aorta exhibited measurably higher RI than those in the cardinal vein. As the dorsal aorta and cardinal vein carry oxygenated and deoxygenated blood, respectively, this contrast is consistent with prior studies demonstrating that hemoglobin’s RI varies with oxygenation state [62], [63], suggesting that the observed RI difference may at least in part reflect differences in blood oxygenation between the two vessels. This offers exciting preliminary evidence that inverse-scattering may be capable of distinguishing arterial from venous blood within scattering vasculature.

For more detailed visualization of specific anatomical features, we examined x-y cross-sections within the dashed white rectangle in **Figure 7(b1)** at depths *z* = 94.76 μm and *z* = 119.48 μm. To achieve clear lateral visualization despite the natural axial curvature of the zebrafish tail, these lateral cross-sections were digitally flattened along the dashed yellow line in **Figure 7(b2)**.

**Figure 7(c1)** shows the x-y cross-section at *z* = 94.76 µm along the embryonic midline. The notochord was prominently visible, characterized by large, vacuolated cells arranged as repeated rounded compartments along the anterior–posterior axis. Individual cell membranes were clearly visible. Dorsal to the notochord, differentiated spinal cord tissue was visible, exhibiting dark, rounded structures consistent with neurons and other neural cell types. The dorsal aorta, clearly visible and indicated by the red dashed line, are the principal vessels running along the ventral surface of the notochord, and groups of red blood cells were clearly localized within its lumen, as expected.

**Figure 7(c2)** shows a zoomed-in view of the posterior region indicated by the yellow dashed rectangle in **Figure 7(c1)**. The neural tube, the early developmental precursor to the spinal cord at this axial level, was clearly visible in this zoomed-in view, and exhibited its characteristic honeycomb-like tessellated appearance arising from tightly packed, proliferating neuroepithelial progenitor cells. The neural tube lumen, a hollow fluid-filled cavity running along the length of the neural tube, is indicated by the blue dashed line, and the floor plate cells adjacent to the notochord are indicated by the green dashed line.

**Figure 7(c3)** shows the x-y cross-section at *z* = 119.48 µm. Distinct myotome bundles forming the segmented trunk musculature were clearly visible with their characteristic V-shaped architecture. Intersegmental vessels branching from the dorsal aorta and extending dorsally between adjacent myotomes were also resolved. **Figure 7(c4)** shows a zoomed-in view of the yellow dashed rectangle in **Figure 7(c3)**, revealing that individual muscle fibers within each myotome bundle can be resolved as elongated, parallel structures oriented along the anterior-posterior axis. Notably, clear striations are visible within individual fibers, arising from the organized sarcomeric architecture. This striated pattern, a defining feature of skeletal muscle, highlights the ability of our approach to resolve subcellular organization in a multicellular organism that is both scattering and absorptive.

## 5. Discussion and Conclusion

In this work, we introduced an inverse-scattering methodology to reconstruct 3D RI and absorptivity in biological samples that exhibit both scattering and absorption. Building on the traditional MSBP framework, we extended the mathematical formulation to explicitly incorporate *complex-valued* RI in the gradient update of the inverse solver. This allowed us to leverage the fact that absorptivity is intrinsically linked to conventional RI as the imaginary component of the more generalized complex-valued RI. While other inverse-scattering approaches have previously demonstrated 3D RI reconstruction in scattering samples, absorption has largely been neglected, either in the reconstruction formulation or in experimental demonstration. To the best of our knowledge, the results presented here constitute the first experimental demonstration of joint 3D reconstruction of both RI and absorptivity in samples that are simultaneously scattering and absorptive.

Interestingly, reformulating the gradient alone was not sufficient to achieve high-quality complex-RI reconstruction in real-world samples. Solving for a complex-valued RI requires solving a highly nonconvex optimization problem over a complex-valued solution space, which is more underdetermined than the corresponding real-valued reconstruction problem. Thus, additional constraints on the search space were essential. In this work, we introduced such constraints by increasing measurement diversity. Specifically, we combined angular-scattering with sample-defocused measurements within the MSBP inverse-scattering framework. Using this strategy, we demonstrated robust reconstruction of volumetric RI and absorptivity across a range of samples including calibration microspheres and biological samples.

There are several promising directions for future research. A natural extension of this work is toward hyperspectral tomography, where illuminating the sample at multiple wavelengths could enable reconstruction of four-dimensional datasets capturing wavelength-dependent volumetric RI and absorptivity. Importantly, spectral dispersion and absorption provide complementary contrast mechanisms that connect tissue structure to composition and physiological state. Access to both could enable powerful label-free differentiation of biological samples that appear morphologically similar yet differ in biochemical makeup, metabolism, or functional activity.

Another exciting direction is to study the challenges introduced by the expanded solution space when reconstructing complex-valued RI compared to real-valued RI. Such an analysis could provide valuable insight on how measurement strategies should be designed to better constrain the inverse problem. For example, our recent companion work has shown that as samples become increasingly thick and scattering, combining angular illumination with defocused measurements is already critical for accurately reconstructing even just real-valued RI [30]. Reconstructing complex-valued RI in similarly thick samples may make the inverse-problem too ill-posed, even after using both angular and defocus measurements. Thus, identifying strategies that provide further constraints is an important direction. Hyperspectral measurements, mentioned above, offer one promising approach. Because RI and absorptivity typically vary smoothly with wavelength, smoothly-varying priors could potentially be leveraged to constrain the inverse problem when a spectral dataset is captured. Another promising direction is to illuminate the sample with structured patterns that are richer and more complex than the angular illumination and sample defocus schemes explored here. Such patterned illuminations could offer greater flexibility in tailoring the measurement diversity to optimally constrain the inverse-scattering problem across different classes of scattering and absorptive samples.

Overall, this work represents an important advance in inverse-scattering methods. By explicitly accounting for absorption, our approach enables accurate reconstruction of both RI and absorptivity in biological tissues, paving the way for more comprehensive, label-free volumetric imaging of biological samples that are both scattering and absorptive.

## Acknowledgment

We thank Jon Tamir and David Fridovich-Keil from UT Austin for insightful discussions on nonconvex optimization. We also thank our lab members for fruitful discussions. Lastly, we acknowledge the Texas Advanced Computing Center (TACC) at The University of Texas at Austin for providing computational resources that have contributed to the research results reported within this paper.

## Data availability

Data underlying the results presented in this paper may be obtained from the authors upon reasonable request.

